# scGREAT: Graph-based regulatory element analysis tool for single-cell multi-omics data

**DOI:** 10.1101/2023.01.27.525916

**Authors:** Chaozhong Liu, Linhua Wang, Zhandong Liu

## Abstract

**Motivation:** With the development in single-cell multi-omics sequencing technology and data integration algorithms, we have entered the single-cell multi-omics era. Current multi-omics analysis algorithms failed to systematically dissect the heterogeneity within the datasets when inferring cis-regulatory events. Thus, there is a need for cis-regulatory element inferring algorithms that considers the cellular heterogeneity.

**Results:** Here, we propose scGREAT, a single-cell multi-omics regulatory state analysis Python package with a rapid graph-based correlation measurement *L*. The graph-based correlation method assigns each cell a local *L* index, pinpointing specific cell groups of certain regulatory states. Such single-cell resolved regulatory state information enables the heterogeneity analysis equipped in the package. Applying scGREAT to the 10X Multiome PBMC dataset, we demonstrated how it could help subcluster cell types, infer regulation-based pseudo-time trajectory, discover feature modules, and find cluster-specific regulatory gene-peak pairs. Besides, we showed that global L index, which is the average of all local L values, is a better replacement for Pearson’s r in ruling out confounding regulatory relationships that are not of research interests.

**Availability:** https://github.com/ChaozhongLiu/scGREAT

## 1. Introduction

Chromatin accessibility refers to the level of physical compaction of chromatin (Minnoye *et al*., 2021). While most of the chromatin regions are densely arranged and hard to be accessed, ~2-3% of total DNA sequences are accessible to transcriptional factors (TFs) that determine the cellular state (Thurman et al., 2012, Klemm et al., 2019). Such accessible DNA regions that regulate the transcription of genes are called cis-regulatory elements (CREs) (Li *et al*., 2015). And it is a well-known marker that reflects the cell state, such as cell development, differentiation, and abnormal disease state. However, it is still one of the major challenges to determine the target genes of CRE activity (Preissl *et al*., 2022) and how CREs’ variations are associated with cellular states.

Recent advances in single-cell technologies and algorithms have enabled the direct link between accessible genome loci and nearby genes. Sequencing techniques like SNARE-seq (Ma *et al*., 2020), SHARE-seq (Chen *et al*., 2019), and 10X Multiome profile the transcriptome and chromatin accessibility simultaneously on the same batch of cells, providing researchers with both modalities’ information; Algorithms like GLUE (Cao and Gao, 2022) and Seurat (Stuart *et al*., 2019), can integrate single-cell RNA-seq and ATAC-seq data, generating data with the two modalities aligned. However, multi-omics analysis tools available now such as Seurat v4 (Hao *et al*., 2021), Liger (Liu *et al*., 2020), MUON (Bredikhin *et al*., 2022), etc., all focus on generating an optimal dimension reduction space utilizing both modalities’ information. However, these tools do not provide downstream analyses, such as CRE inference, after integrating the two modalities.

One common method to infer CREs is the co-accessibility measurement (Preissl *et al*., 2022). For example, Cicero utilizes scATAC-seq data to connect distal regulatory elements with target genes by computing a covariance matrix of accessible sites (Pliner *et al*., 2018). ArchR (Granja *et al*., 2021), another popular scATAC-seq analysis software, applies Pearson correlation (Benesty *et al*., 2009) to infer the links between scATAC-seq and scRNA-seq data. Such correlation-based methods could fairly detect CREs in homogenous population of cells. However, since cellular heterogeneity is not modeled in their approaches, these methods could miss regulatory relations that are specific to subpopulations of cells. These cell subpopulations could represent different cell types (Grün *et al*., 2015), development stages (Velten *et al*., 2017), and more. Although diverse bioinformatics methods are available to either annotate cell types (Clarke *et al*., 2021) or infer cell development trajectory (Saelens *et al*., 2019), the origin of heterogeneity is more complicated and any attempt to divide cells into subpopulations is a simplification and information loss. Thus, a systematic detection of regulatory heterogeneity without an assumption on heterogeneity origin is needed.

In geographical studies, researchers utilize measurement specially designed for spatial data to dissect the correlations between features. This kind of metrics considers the spatial pattern in correlation, and provides a correlation measurement for each location within its neighborhood, which is called the local correlation value. Such local correlation value represents the heterogeneity of feature correlation among all the locations, and the sum of all local values is the general correlation trend, or the global correlation. We see the resemblance of spatial correlation between the geographical and single-cell multi-omics studies since multi-omics data could be represented in a low-dimensional space and constructed as a k-nearest neighboring (KNN) graph. Thus, it is feasible to develop a similar global and local correlation measurement to dissect the general correlation and cellular regulatory heterogeneity.

Here, we adapted the geographical spatial correlation measure (Lee, 2001) to single-cell studies and developed the Python package – scGREAT (single-cell Graph-based Regulatory Element Analysis Toolkit) to infer single-cell resolved regulatory states using local *L*-index values. First, we will show how the local *L* matrix has enabled the comprehensive study of heterogeneity in the 10X Multiome PBMC dataset, including sub-clustering cell types, inferring regulation-based pseudo-time trajectory, discovering feature modules, and finding cluster-unique regulatory relationships. Then we will demonstrate why our global *L* index, which averages local *L*-index values across all cells, is a better replacement for Pearson’s *r* in studying the general gene-peak regulatory trend by ruling out the confounding gene-peak pairs that are not of research interests.

## 2. Methods

### 2.1 Single-cell multi-omics data preprocessing

Three datasets from common multiome techniques, including 10X Multiome, SHARE-seq, and SNARE-seq, have been employed to test the feasibility of our software. 10X Multiome Peripheral Blood Mononuclear Cells (PBMC) data were processed using Seurat v4 by following the Seurat tutorial at https://satijalab.org/seurat/articles/weighted_nearest_neighbor_analysis.html (Hao *et al*., 2021). Then the scRNA-seq and scATAC-seq count matrices, nearest neighbor graph and metadata were exported and loaded in Python to create the final AnnData object (Virshup *et al*., 2021) as the input of our package.

SHARE-seq mouse skin and SNARE-seq mouse brain datasets were downloaded at GEO by GSE140203 and GSE126074. The raw count matrices were loaded and preprocessed using Scanpy (Wolf *et al*., 2018) following the standard preprocessing pipeline (https://scanpy-tutorials.readthedocs.io/en/latest/pbmc3k.html). The final AnnData objects were saved as the input for our package. All codes are available in the GitHub repository.

### 2.2 Graph-based correlation index *L*

The graph-based correlation measurement is the core of our package. The correlation index *L* between gene vector 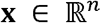 and peak vector 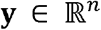 combines the normal correlation score and graph dependence values between the two features. It is defined as:

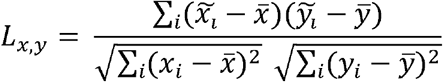

where *i* ∈ {1,.., *n*} represents cell index, 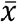 and 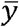 are the numeric mean values of **x** and **y**, 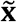 and 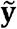 are the spatial lag values which are composed of weighted averages of cell neighbors. Spatial lag of *x_i_* is defined as:

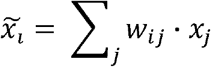

Where *j* is index of connected cell with *i* in the graph, and *w_ij_* is their connectivity weight. Here, we take the K-nearest neighbor graph derived from the dimensional reduction results as the connectivity matrix 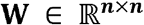, where *n* indicates the cell number.

One major challenge of implementing such approach for single-cell data is the enormous number of gene-peak pairs, which is computational infeasible. To speed up the computation and allow scalability, we used a vectorized implementation. Suppose **W** is the row-standardized (row sums are all 1) connectivity matrix, **Z** is an z-scored 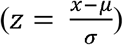 form of 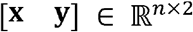, **L** between **x** and **y** can be calculated with:

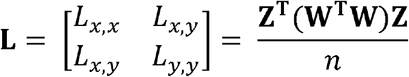

When implementing the calculation in Python, we vectorized the formula again to include all *p* gene-peak pairs at once:

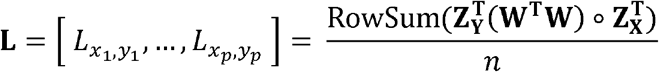

where **Z_X_** and **Z_Y_** are z-scored matrices of the RNA-seq log-transformed data 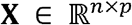, and scATAC-seq log-transformed data 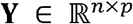.

The local value of *L*, associated with each cell, indicates each cell’s relative contribution to the global *L*. Local *L* index of cell *i* between gene **x** and peak **y** is defined as:

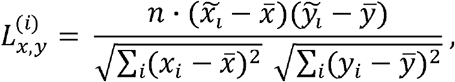

allowing it to capture a cell’s association within its neighborhood.

The vectorized implementation of the local *L* matrix ∈ 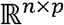 can be calculated with:

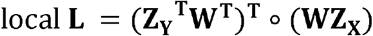

To speed up all the graph (spatial) dependence measurement, we also vectorized the implementation of Moran’s *I* in the package. Details are discussed in the supplementary materials (Package Details 2.4). We evaluated the computational time under the constrain of 16 Gb memory and eight threads. In summary, the graph-based measurements in scGREAT are much faster than the original implementation in geographical package *esda* (Rey and Anselin, 2007) (Figure 1B).

**Figure 1.**
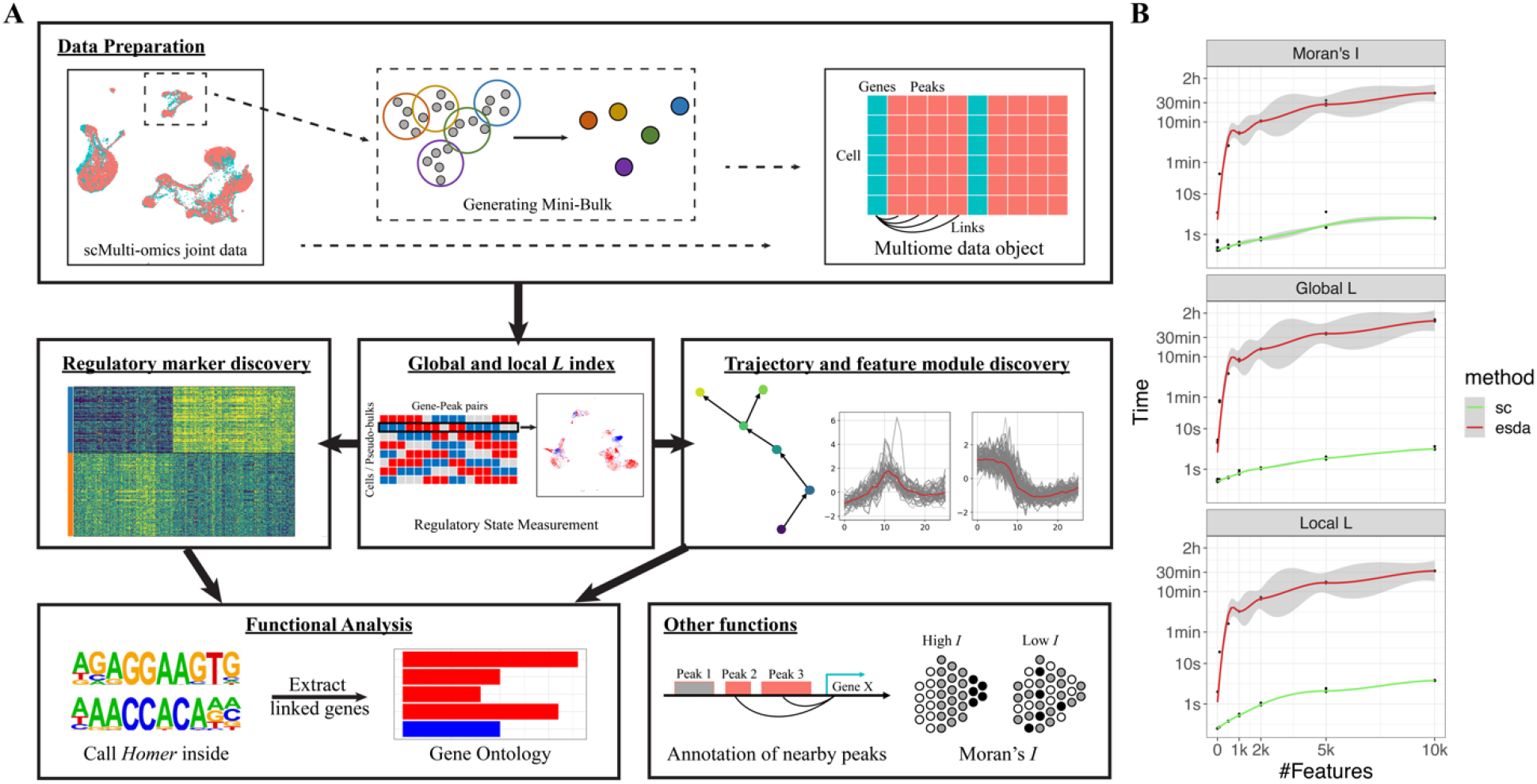
scGREAT overview. (A) Analysis pipeline of scGREAT. scGREAT receives as input the integrated scRNA-seq and scATAC-seq data. Users can choose to generate pseudo-bulk data or skip this step and use the original single-cell data. After linking genes with nearby peaks, scGREAT will preprocess the raw data and generate the final multiome data object for further analysis. Next, before all other analyses, the *L* index will be computed with user-chosen parameters. With the global *L* value and local *L* matrix, users can perform labeled analysis (regulatory marker discovery) and unlabeled analysis (sub-clustering, trajectory, feature module discovery), followed by functional analysis to help explain *Homer* output results. (B) Computation time comparison between scGREAT and *esda*. Under the limitation of 16 Gb memory and eight threads, the time consumed to calculate Moran’s *I*, global *L*, and local *L*, was estimated with different feature numbers.

To quantify the statistical significance of L index, we implemented a permutation test. Specifically, the paired vectors were shuffled together *n* times to get a reference distribution of the *L* index. Then the z-score of the *L* index under the reference distribution is calculated and taken as the simulated p-value. But in single-cell studies, data are usually sparse (Lähnemann *et al*., 2020) and of large sample size (cell number), which results in all significant p-values during initial experiments. Here, we introduce a conditional permutation process. In each permutation, we only shuffle a certain percentage (default 10%) of values and permutate among non-zero and zeros values (see pseudo-code in supplementary material). After permutation, the same process mentioned above will be done to get the final p-value as the significance level.

### 2.3 Implementation of all package functions

**Figure 1A** shows the analysis pipeline and all functions in scGREAT. Modules will be briefly described here, and all details can be found in the supplementary materials.

#### Data preparation

The input of scGREAT is scRNA-seq and scATAC-seq data that have already been aligned. It can be naturally paired from multiome sequencing techniques or aligned by integration algorithms. Users can use single-cell data for the analyses or generate pseudo-bulk data using scGREAT functions. After annotating genes with their nearby peaks, the final multiome AnnData object that combines transcriptome and chromatin accessibility data will be generated for the other modules.

#### Graph-based measurements

The graph-based measurements are the core of scGREAT, including Moran’s *I*, global *L* index, and local *L* matrix. Users just need to run a one-line code to perform each measurement.

#### Unlabeled analysis

When users are interested in one or few clusters of cells or when no label information is available, unlabeled analysis can be done to perform sub-clustering and trajectory analysis based on the regulatory state measured in core functions. With pseudo-time inferred, self-organized map (SOM) algorithm was implemented to discover feature modules.

#### Labeled analysis

When users have already annotated the data with biological groups, a statistical test can be done to discover all the differentially regulated pairs among all the groups using the local *L* matrix calculated.

#### Motif Enrichment Analysis

With a list of peaks derived from any analysis, we implemented functions to prepare input and call *Homer* (Heinz *et al*., 2010) for motif enrichment analysis. There is also a function helping extract peaks and related information from Homer results with users’ motifs of interest.

#### Visualization

To help users understand the results and generate publishable plots, we developed visualization functions for each step and module, including pseudo-bulk quality summary, local *L* index heatmap, feature plot in UMAP, volcano plot after statistical test, and feature module plot after SOM implementation.

## 3. Results

### 3.1 Pseudo-bulk data shows higher correlations between gene and promoter regions

The first step for the single-cell multi-omics regulatory element analysis is data preparation. In this step, generating pseudo-bulk data is recommended. Here, we measured the correlation between gene expression and promoter accessibility by the graph-based *L* index and Pearson’s *r* in three datasets generated from common multiome techniques, including 10X Multiome, SHARE-seq, and SNARE-seq. Both methods showed a higher level of correlation in pseudo-bulk data compared with the single-cell data (Figure 2). We believe this is due to the single-cell data sparsity (Lähnemann *et al*., 2020) that makes the data nosier than bulk ones. When generating pseudo-bulks averaging the values of several cells within a local neighborhood, the data is less noisy, and thus the results have a better correlation signal. With this observation, in the package, we provide the function to generate pseudo-bulk data (see supplementary method for details) for cis-regulatory element discovering. The following results are all based on the pseudo-bulk data generated by our package.

**Figure 2.**
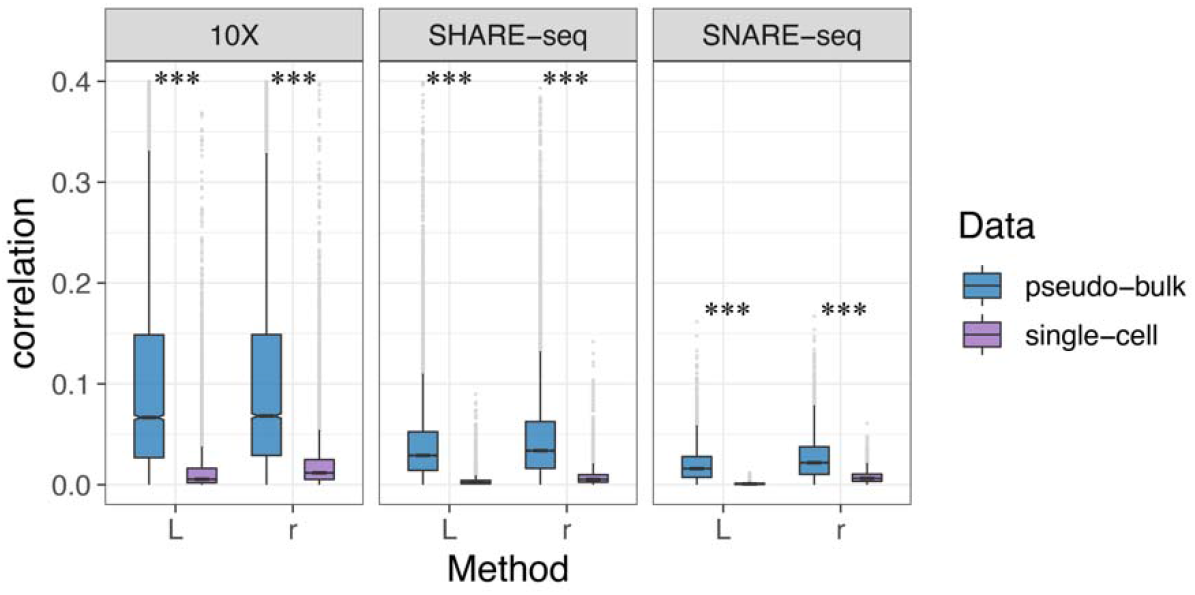
The correlation between genes and their nearest upstream peaks by *r* and *L* in pseudo-bulk and single-cell data. Three datasets were applied, including 10X Multiome PBMC, SHARE-seq mouse skin, and SNARE-seq mouse brain, to compare pseudo-bulk and single-cell data. Paired t-test was done for significance.

### 3.2 Local *L* index as an indicator for cell regulatory state

The local *L* index indicates the cell’s gene-peak correlation strength (magnitude) and direction (sign) within its neighborhood. For each gene-peak pair, local *L* is calculated for each cell to measure single-cell resolved regulatory states. By doing this for all cells and all gene-peak pairs, we generated a new layer of information representing the regulatory state besides the transcriptome and chromatin accessibility matrix. Here, we will demonstrate how local *L* index and the cell regulatory state matrix can help study the regulatory heterogeneity in the well-annotated 10X Multiome Peripheral Blood Mononuclear Cell (PBMC) dataset.

Supplementary Figure 1 shows examples of such regulatory states between genes and peaks. RORA and chr15:60649315-60650653 in CD4+ T cells, for example, are not highly correlated (Pearson’s *r* = 0.404). However, the regulatory state defined by the local *L* shows that the correlation is high in CD4+ T naïve cells and CD4+ TEM cells, but not in the majority of CD4+ TCM cells. And this heterogeneity was overlooked by Pearson correlation. RORA is known to function in T cell differentiation (Solt *et al*., 2011). The correlated peak lies upstream of isoform b and c of RORA, and the regulatory state changes could indicate the alternative splicing and different functions in cell differentiation. Thus, within-cluster heterogeneity could also be important in understanding the cellular process besides the global correlation trend measured with Pearson’s *r*.

With this cell-specific regulatory information encoded in the cells-by-regulatory pairs matrix, scGREAT enables cluster-specific CRE analysis. Under the assumption that regulatory states indicate cell state, such as cell differentiation, the local *L* matrix is utilized for sub-clustering and trajectory analysis in scGREAT similar to scRNA-seq analysis. Here, we used the same 10X Multiome CD4+ T cells as an example. After sub-clustering in the pseudo-bulk data, scGREAT mapped the labels back to the single-cell data, shown in Figure 3A left. The regulatory state-derived clusters split the three manually annotated cell types into more subtypes. Using the same KNN graph constructed in the sub-clustering process, trajectory analysis was performed with functions in scGREAT utilizing diffusion pseudo-time (Haghverdi *et al*., 2016) implemented by Scanpy (Wolf *et al*., 2018), and the pseudo-time labels were transferred back to single-cell data (Figure 3A middle). While the trajectory agrees with the pseudo-time inferred using scRNA-seq data only (Figure 3A right), the regulatory state-based results were smoother and better separated cell types CD4+ TCM and CD4+ TEM as shown in the density plots (Figure 3B).

**Figure 3.**
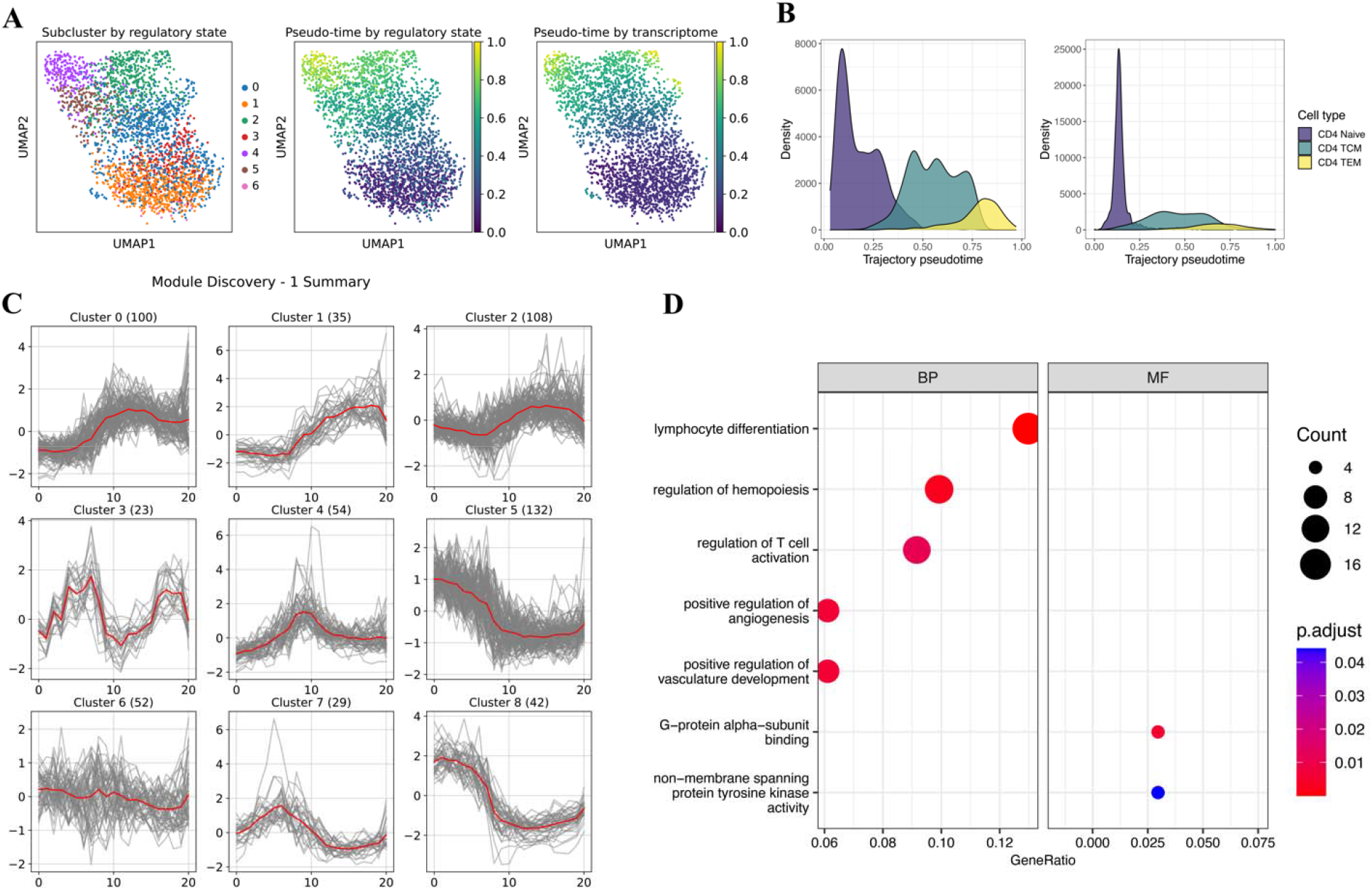
Heterogeneity analysis on CD4+ T cells. (A) UMAP labeled by subclusters (left), trajectory pseudo-time (middle) derived from regulatory state, trajectory pseudo-time (right) derived from the transcriptome. (B) Density plots showing the pseudo-time distribution in regulatory state and transcriptome-derived trajectory, grouped by cell sub-types. (C) Visualization of the self-organizing map (SOM) optimization results. It demonstrates the features modules found by SOM. (D) Gene Ontology enrichment results of genes in clusters 5 and 8. Size represents the number of genes; color represents the adjusted p-value.

To dissect the trajectory pattern, we implemented the self-organizing map (SOM) (Vettigli, 2018; Kohonen, 1990) in our package for feature module discovery such that features with similar temporal patterns will be clustered together. The SOM algorithm is an unsupervised artificial neural network that groups all observations in a low-dimensional representation while preserving the topological structure of the data. Here, we take genes or peaks as observations and the averaged expression levels within each pseudo-time bins as features, and clustered the genes or peaks by SOM. Users can run the function multiple times to have the optimized results by changing the SOM shape (how many modules are needed and what is the similarity among all modules), learning rate, number of iterations, and regulation power sigma. After optimizing the SOM, scGREAT can visualize the results to give users an intuitive understanding how genes or peaks are changing along the pseudo-timeline. In our example of CD4+ T cells analysis, the self-organizing map grouped all the genes into nine modules with different patterns (Figure 3C). Module 5 and module 8 have similar trends in which high expression in CD4+ Naïve T cells dropped in CD4+ TCM and raised slightly in CD4+ TEM cells. We performed Gene Ontology enrichment analysis to determine the function of the two gene modules, shown in Figure 3D. The functions discovered by GO analysis are consistent with the cellular process of the inferred trajectory and CD4+ T cells, including lymphocyte differentiation and T cell activation. In our regulatory state analysis, genes and peaks are always linked. Following the detection of gene-linked peaks in module 5 and 8, we performed motif enrichment analysis with *Homer* (Heinz *et al*., 2010). All top enriched motifs are from the ETS transcription factors family binding domain and RUNX1 binding domain. Such results are consistent with the previous findings that ETS family of transcription factors are related to lymphoid differentiation (Russell and Garrett-Sinha, 2010). Moreover, RUNX1 functions in the development of normal hematopoiesis (Fujimoto *et al*., 2007), and controls the anergy and suppressive function of regulatory T-cells (Ono *et al*., 2007).

### 3.3 Regulatory marker discovery based on the local *L*

With the regulatory state matrix from local *L* and cluster labels, we implemented the t-test for differentially correlated gene-peak pairs in the same way as DEGs (Differentially Expressed Genes) and DARs (Differentially Accessible Regions) analysis. If a certain gene-peak pair is differentially correlated in one cluster compared with all other clusters, it is considered a potential regulatory marker worth further studying. In contrast, Pearson correlation which doesn’t have such a local representation, cannot achieve the same aim with only a single value. Here, we used functions (*FindAllMarkers* and *FindMarkers*) in scGREAT to find all regulatory markers in peripheral blood mononuclear cell types (Supplementary Figure 2A). With scGREAT, regulatory state differences can be visualized in a heatmap with cell labels annotated (Supplementary Figure 2B). Next, scGREAT functions can help select significant markers of each cluster by the mean difference, adjusted p-value, and feature sparsity, then visualize the differential regulatory pairs with the volcano plot (Supplementary Figure 2C). If a pair is of great interest, the local *L* index in pseudo-bulk data can also be mapped back to the original single-cell data (Supplementary Figure 2D, Package Details 1.5).

With this regulatory marker discovery method, we studied the regulatory changes in B cell development and CD4+ T cell differentiation (Figure 4A, B). After the markers have been found, we extracted the peaks and performed motif enrichment analysis. Besides RUNX1 binding domain mentioned previously in T cell differentiation, the PU.1 binding motif was enriched in B cell development. This transcription factor is encoded by the Spi1 gene, binds with the PU-box, and activates gene expression during myeloid and B-lymphoid cell development (le Coz *et al*., 2021). Genes linked with these peaks are potentially the targets of the transcription factors. So, we collected the genes potentially regulated by PU.1 and RUNX1, and performed Gene Ontology enrichment analysis (Figure 4C, D). During B cell development, regulation of antigen receptor-mediated signaling pathway has changed, as well as the immunoglobulin subunit Fc receptor-mediated stimulatory signaling pathway. During naïve T cell differentiation, antigen processing and presentation process has changed; lymphocyte anergy functions have also changed, indicating the activation of CD4+ T cell from Central memory cells to effector memory cells. All these analyses have validated the markers discovered by our method.

**Figure 4.**
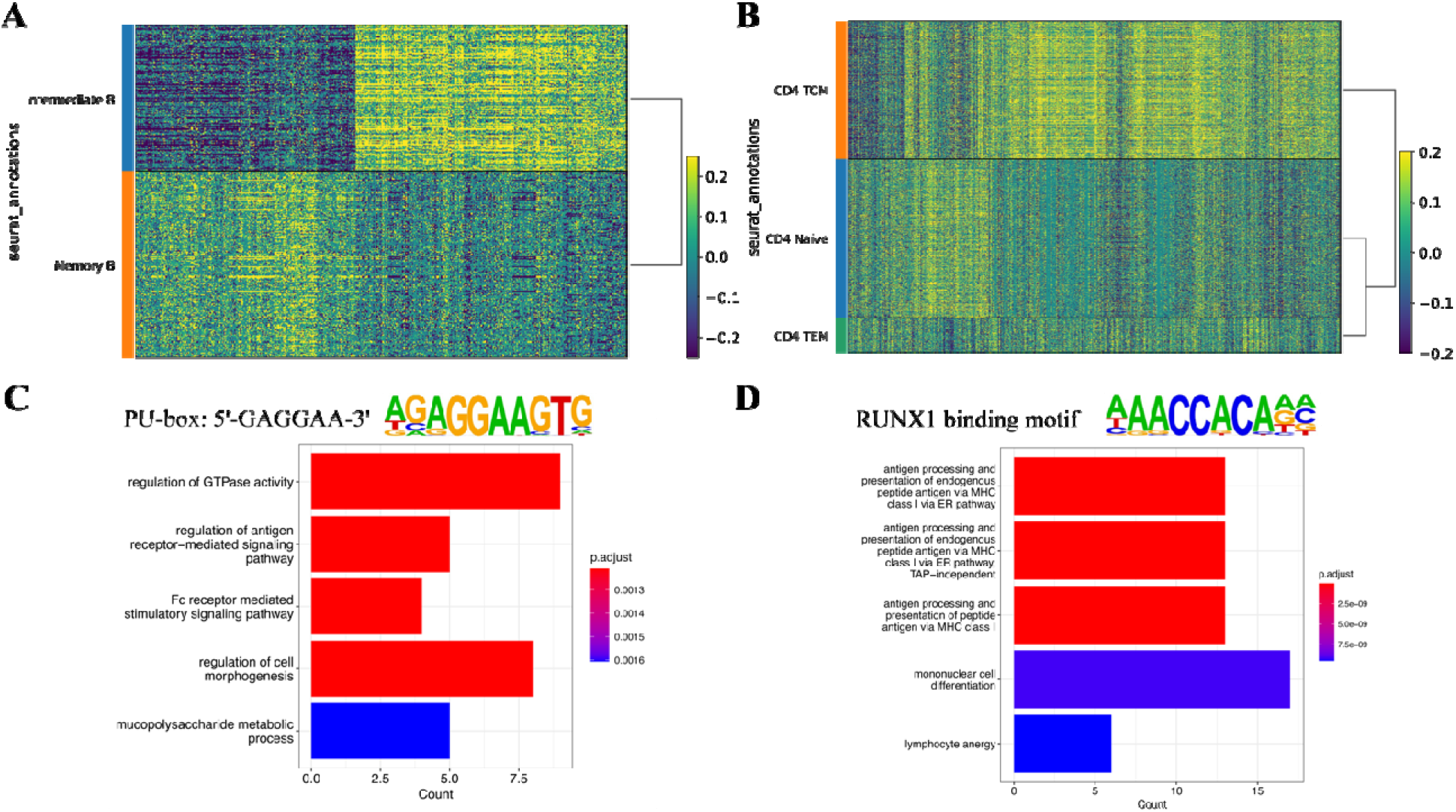
Markers discovery on 10X Multiome PBMC dataset. (A) Heatmap showing the gene-peak correlations in pseudo-bulk data of B cell clusters. Each row is a pseudo-bulk, each colum**n** is a gene-peak pair, and color indicates the correlation level. (B) Heatmap showing the gene-pe**ak** correlations in pseudo-bulk data of CD4+ T cell clusters. (C) Differentially correlated peaks in **B** cell clusters were enriched in PU-box binding motif. Genes linked with the peaks were extract**ed** for Gene Ontology enrichment analysis to discover the functional changes. (D) Differentia**lly** correlated peaks in CD4+ T cell clusters were enriched in the RUNX1 binding motif. Gen**es** linked with the peaks were extracted for Gene Ontology enrichment analysis to discover t**he** functional changes.

### 3.4 The global *L* index is a better replacement for Pearson’s *r*

The global *L* index, which averages local *L*-index values across all cells, serves the same role as Pearson’s correlation to show the general correlation trend in the data. Both Pearson’s and our graph-based (*L*) correlation are variants of Mantel’s general cross-product association measure (Mantel, 1967). But the graph-based correlation takes the single-cell dataset as a K-nearest neighbor (KNN) graph and cares about the neighborhood patterns, which varies from Pearson correlation (see Method 2.2 for details).

When measuring the general correlation trend (no consideration in heterogeneity) within the dataset, *L* index and Pearson’s *r* are consistent. This pattern was observed in all three datasets (Figure 5A), with statistically significant high correlations. This consistency was still there when analyzing a single cluster rather than the whole dataset (Supplementary Figure 3A).

**Figure 5.**
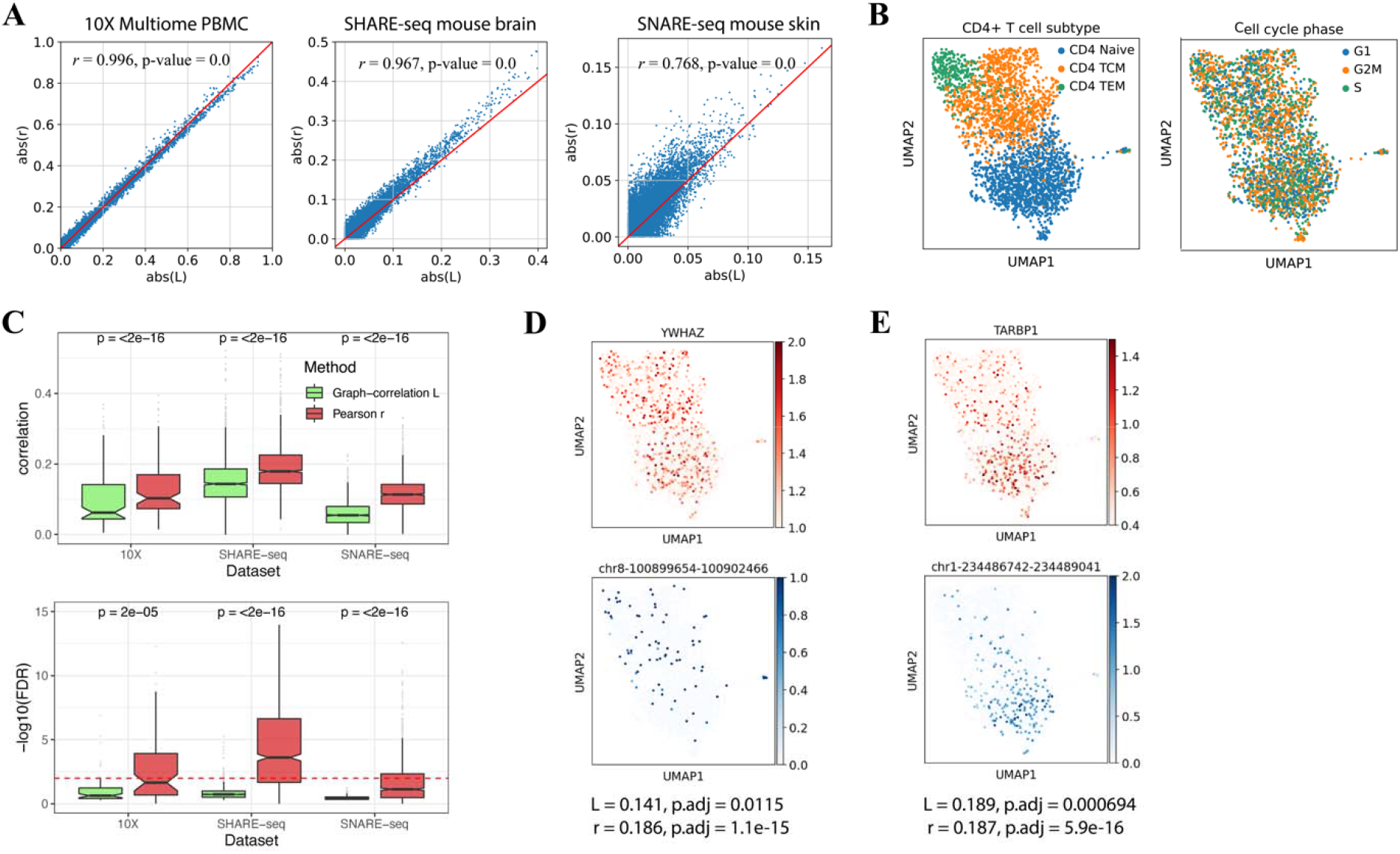
Comparison between Pearson’s coefficient *r* and global *L* index. (A) The consistency between *L* and *r* in studying the general trend of regulatory relationships. The scatter plots showed all gene-peak pairs correlation levels measured with *r* (y-axis) and *L* (x-axis). The consistency between *r* and *L* was tested significantly by Pearson correlation (results shown in the plots). (B) UMAP visualization of CD4+ T cells labeled by T cell subtype (left) and cell cycle phase (right). (C) Box plots showing the correlation level (top) and significance test FDR (bottom) measured by *L* and *r*, between cell cycle genes and their nearby peaks. (D) UMAP visualization of cell cycle gene YWHAZ (top), and its cis-regulatory element chr8-100899654-100902466 (bottom), (E) immune-related gene TARBP1 (top), and its cis-regulatory element chr1-234486742-234489041 (bottom). The two pairs have similar Pearson correlations but because of the pattern in data structure, their *L* index and significant test differ.

Beyond the consistency, only the global *L* index can rule out potentially confounding regulatory pairs that are not of research interests, which is an advantage of our method over Pearson’s r. To demonstrate this observation, we took the CD4+ T cell cluster from 10X Multiome data as an example. In Figure 5B, we visualized the data by UMAP and labeled the cells with their cell cycle phases and differentiation stages. It is clearly shown that cell cycle phases are mixing in the UMAP while the differentiation stages are separated off. Thus, we believe the CD4+ T cell data is about T cell differentiation and activation. And the regulatory pairs between cell cycle genes and their nearby cis-regulatory elements, as one of many examples, are confounding pairs to the research. To infer the regulatory relationships in T cell differentiation, we would like the correlation method to exclude such confounding pairs.

Here, by taking cell cycle genes and their nearest peaks (most likely to be promoter regions) as confounding examples, we compared the behavior of *L* index and Pearson’s *r* in CD4+ T cells from 10X Multiome data, cluster 7 from SHARE-seq data, and cluster 2 from SNARE-seq data. Figure 5C shows the correlation (top) and significance test (bottom) between cell cycle genes and their nearest upstream peaks. The *L* index tends to have smaller values than Pearson’s *r* and distinguished the confounding pairs through significant test. We think it is because such cell cycle-related gene expression and peak accessibility are uniformly distributed in the data (examples in Supplementary Figure 4A). Though they correlate well, there are low and high values mixing within each cell neighborhood. Such discordance in the neighborhoods will influence the graph-based *L* index measurement and significance test. On the contrary, the immune-function related gene TARBP1 has a higher *L* index with its nearby peak compared to cell cycle gene YWHAZ and its nearby peak, though the two pairs have the same Pearson’s correlation *r* (Figure 5D, E). By taking the neighborhood consistency into the correlation measurement, the global *L* index ruled out such confounding regulatory pair. However, Pearson correlation doesn’t know how cells are distributed in the data space and cannot achieve the same effect.

In practice, this behavior of our correlation measurement could enhance the specificity for downstream analysis. For example, when gene-peak pairs are ranked by correlation and taken for Gene Set Enrichment Analysis (GSEA) (Subramanian et al., 2005), ruling out such confounding pairs will make the molecular process and pathways of research interest stand out from the pool. Here, we compared the GSEA results from L index-based and Pearson’s r-based gene ranks. Because of the difference in ranks, only 96 out of 8995 gene sets were significantly enriched from Pearson’s r-based ranks, while 355 were discovered in L index-based ranks (adjusted p-value < 0.01). Then, we collected all T cells related sets and compared the results. In general, T cell-related gene sets rank higher from L-based correlation (Supplementary Figure 4C). Among the 15 significantly enriched T cell sets in L index-based results, only 4 can be discovered with Pearson’s r-based ranks (Supplementary Figure 4D).

## 4. Conclusion and Discussion

scGREAT is a novel single-cell multi-omics analysis toolkit with a rapid graph-based correlation measurement and related analysis functions. Compared with other correlation methods like Pearson correlation, the graph-based correlation method cares about spatial dependence and poses a specially designed significance test for single-cell data. It thus enabled the filtering of confounding regulatory pairs out of research interests and kept only regulatory pairs with patterns consistent with the main biological variance. Besides, with the local *L* index from the correlation method, scGREAT can generate the regulatory state matrix, which is a new layer of information. With the graph-based correlation scores, scGREAT filled the gap in multi-omics regulatory analysis by enabling labeled and unlabeled analysis, functional annotation, and visualization.

Pearson correlation, our graph-based correlation, and any other variants of Mantel’s general cross-product association measure (Mantel, 1967), are affected by dropouts in single-cell data. After scaling the data with zero means and unit variances, dropout values will turn to negatives from zero. These negative but meaningless values will somewhat bias the final correlation results. To deal with this problem, implementations in single-cell scenarios need to pay special attention to feature sparsity and dropouts in all processes. In scGREAT, first, we offer users pseudo-bulk analysis options to decrease the dropout rate. Second, we keep the global *L* index with no modification, but offer users feature sparsity information to measure the influence of dropouts on final values quantitatively. Then for the local *L* index or the regulatory state matrix, scGREAT will zero the values when it is derived from any of the two features with dropout values, making the final matrix similar to any single-cell data with dropouts. Implementing this process is beneficial for both labeled and unlabeled analysis to avoid misleading results. Nevertheless, the future direction in measuring multi-omics regulatory relationships could be to avoid the effect of dropouts.

The groundbreaking Spatial-omics techniques have arisen in recent years (Asp *et al*., 2020; Marx, 2021; Deng *et al*., 2022). These techniques profile omics data and map them back to the two- and three-dimensional geography of tissues, allowing researchers to study cell-cell communication and tissue organization (Ståhl *et al*., 2016; Moffitt *et al*., 2016; Deng *et al*., 2022). Our *L* index is a good fit for the gene co-expression analysis in spatial omics data. The *L* index considers spatial patterns in correlation measurements, and the local *L* index provides the correlation pattern in the geography space. To fit scGREAT into spatial data analysis, we need to replace the connectivity matrix with the spatial connections-based matrix, which is feasible. Thus, our future direction will be how to apply scGREAT to spatial transcriptomics spatial co-expression analysis and multi-omics spatial correlation analysis.

## Data and Code Availability

The data underlying this article are publicly available (10X Multiome PBMC: https://support.10xgenomics.com/single-cell-multiome-atac-gex/datasets/1.0.0/pbmc_granulocyte_sorted_10k, SHARE-seq mouse skin: GSE140203, and SNARE-seq mouse: GSE126074).

All codes to replicate the results are available at the GitHub repository.

## Funding

Research reported in this publication was supported by the Eunice Kennedy Shriver National Institute of Child Health & Human Development of the National Institutes of Health under Award Number P50HD103555 for use of the Bioinformatics Core facilities. The content is solely the responsibility of the authors and does not necessarily represent the official views of the National Institutes of Health. ZL, CL and LW are also partially supported by the Chao Endowment and the Huffington Foundation. The funders had no role in study design, data collection and analysis, decision to publish, or preparation of the manuscript.

## Acknowledgements

The authors would like to thank members of the Liu lab for their suggestions and discussion.

## Notes

https://github.com/ChaozhongLiu/scGREAT

### Competing Interest Statement

The authors have declared no competing interest.

