## Supplementary Materials for "scGREAT: Graph-based regulatory element analysis tool for single-cell multi-omics data"

### Supplementary Methods

#### 1. Estimation of computation time

We estimated the time consumption for calculating Moran's *I*, global *L*, and local *L* index because the three are commonly required for analyzing single-cell data. To make a fair comparison between scGREAT and *esda*, we limited the computational capacity to 8 threads and 16 Gb of memory. This is done by setting the number of threads with *os.environ ['OPENBLAS_NUM_THREADS']* and set the *max_RAM* parameter of the three functions to 16. No memory limitation was set for esda since it never used more than 16 Gb memory during the experiment.

To test how time consumption changes along with the number of features, we used the 10X Multiome PBMC dataset with 5000 pseudo-bulks and selected feature numbers to be 1, 10, 100, 500, 1000, 2000, 5000, and 10000. Time was estimated with the *time* package in *Python*.

#### 2. Cell cycle related analysis

The cell cycle genes were selected from KEGG cell cycle pathways (human, mouse) to compare the correlation level of the global *L* index and Pearson's *r* in Figure 2D.

Human cell cycle phase markers were selected from the tinyatlas GitHub repository to predict the cell cycle phase of all CD4+ T cells in Figure 2C. With the list of markers, we used the *score_genes_cell_cycle* function in *Scanpy* to predict the phases with average marker expression levels.

#### 3. Functional analysis

Gene Ontology enrichment analysis and visualization were done using the *clusterProfiler* library in *R*. GSEA (Gene Set Enrichment Analysis) was done using the *fgsea* library in *R*. We selected curated gene sets (C2) and ontology gene sets (C5) in the enrichment. Motif enrichment analysis was done using *Homer* called from the scGREAT package. All related results were provided in supplementary materials.

**4. Statistical test**

All paired t-test results above the box plots were calculated by *stat_compare_means* from the *ggpubr* library in *R*. Pearson correlation coefficient and p values were calculated by *scipy.stats.pearsonr* in *Python*.

#### Package Details

In this section, details about the package are explained in the order of the pipeline. Along with the explanation, usage detail and recommendation will also be discussed.

**1. Data preparation**

**1.1 Input data**

The expected input of the package is integrated single-cell multi-omics data. It can be from the single-cell multiome sequencing techniques or the integration results of algorithms like Seurat and GLUE. Specifically, the package needs the AnnData object with the following information:

1. Raw count matrix of gene expression and chromatin accessibility. Cells in the two matrices should be ordered to reflect the one-to-one correspondence between omics data. So, cell numbers are the same.
2. Joint embedding space. This is the embedding space where all cells from ONLY one omics data are projected. It is needed to construct the nearest neighbor map. Examples: PCA from the Seurat analysis of the 10X Multiome data, PCA from the analysis of only scRNA-seq data, integrated joint embedding space from GLUE including only scRNA-seq cells, etc.

3. Metadata (optional). Include the metadata information like sample ID, and cluster annotation for further analysis.

**1.2 Data preparation steps**

i. *nn_map* function to generate the nearest neighbor map
ii. *add_group_sparsity* to calculate the feature sparsity in the whole data or clusters (optional), because sparsity is needed to filter correlated gene peak pairs found.
iii. (optional) generate the pseudo-bulk data with *PsedoBulk*. Users can provide the *group_name* parameter indicating the metadata column name to generate all pseudo-bulks within each cluster label.
iv. Preprocess the RNA-seq data with *RNA_preprocessing_in_one*. It will filter genes with sparsity higher than the cutoff, normalize and log-transform the data, determine highly-variable genes if not provided by the user, perform PCA, construct K nearest neighbor graph, and perform UMAP dimensional reduction.
v. Preprocess the ATAC-seq data with *ATAC_preprocessing_in_one*. It will filter peaks with sparsity higher than the cutoff, normalize and log-transform the data, and calculate all peaks' mean and standard deviation.
vi. Link genes with nearby peaks within the range defined by users by providing parameters *upstream* and *downstream* in *peaks_within_distance* function.
vii. Generate the multiome AnnData object with *multiome_data*. The final multiome object contains a log-transformed matrix of genes and peaks, the K nearest graph for *L* index calculation, and links between peaks and genes.

**1.3 Single-cell multi-omics pseudo-bulk data generation**

As discussed in the results, pseudo-bulk can reduce the noise in single-cell data. Here we implemented the pseudo-bulk generation similar to Cicero [ref] and embedded it as an easy-to-run function in the package. With the integrated single-cell multi-omics data, the package will calculate the nearest neighbor map among all the cells using the embedding space that users provided. Then the package will repeatedly and randomly select neighborhoods and sum up their omics data, until the pseudo-bulk number reaches either the user-defined maximum or the number of cells. Besides, the function will skip neighborhoods that have more than 90% cell overlap with previous pseudo-bulks.

**1.4 Fast annotation of genes with nearby peaks**

Before generating the multiome data object, we need to define the link between genes and peaks. Usually, the link is a user-provided genome range upstream and downstream of the gene body. Peaks within the range potentially interact with the gene expression process and are thus linked. In the package, we provided a function called "*peaks_within_distance*" to perform the task utilizing *Pandas*-embedded parallel programming. It judges whether the mid-point of peaks locates within the range defined near a particular gene, and provides information, including the distance to the transcription starting site, distance to the transcription ending site, whether it's in the promoter region, and whether it's in the gene body. By default, it will remove the link between the gene and the peak within another gene body. This can be disabled by setting *no_intersect* to False.

**1.5 Mapping results back from pseudo-bulk to single cell**

When all analysis is done in pseudo-bulk data, the results like *L* index, or trajectory pseudo-time, are likely needed by the single-cell data. To achieve this, we implemented the function *map_back_sc* to transfer the numeric or categorical results back to each cell, by taking the average (numeric data) or max votes (categorical data) of all the pseudo-bulks containing that cell.

#### 2. Calculate the global *L* index and local *L* matrix

The global *L* index and the local *L* matrix is the basis of all analysis embedded in the pipeline.

##### 2.1 Global *L* index

With the multiome data object created, users can calculate the global *L* index to study the regulatory homogeneity by calling *Global_L*. The results will be saved in the AnnData.uns[‘peaks_nearby’] containing the *L* index, p-value, and FDR if the *permutations* parameter is not 0. Besides, Pairs can be filtered by feature sparsity by using the *percent* parameter.

##### 2.2 Pseudo-code implementation of the significance test for *L* index

**Algorithm** Significance test for global *L* index

```
for x ∈ ℝ^n, y ∈ ℝ^n
L_ref = [ ] (Reference distribution of L_{x,y})
repeat n times
    x', y' = shuffle 10% x, y together
    append L_{x'y'} to L_ref
return p = (L_{x,y} − L̄_ref) / std(L_ref)
```

$$p = \frac{L_{x,y} - \bar{L}_{ref}}{std(L_{ref})}$$

##### 2.3 Local *L* matrix

To speed up the measurement, the function *Local_L* will first filter pairs by the gene and peak Moran's *I* value cutoff provided by the user. Moran's *I* can be calculated with the *Morans_I* function. If Moran's *I* is not found within the *Local_L* function, it will automatically run *Morans_I* first. After calculating the local *L* index matrix, users have the option to remove the effect of dropout values. Specifically, if the gene-peak local *L* value is from cells that don't express the gene or have the peak opened (i.e., value is 0, and most likely to be dropout), the *L* value will be changed to 0. According to our experiment, this option is recommended and defaulted in the *Local_L* function. But users can also set the *dropout_rm* parameter to False to disable the behavior.

If the local *L* matrix is for cluster regulatory marker discovery, then the Local L matrix will be calculated per cluster within the *FindAllMarkers* or *FindMarkers* function. Users don't need to call the *Local_L* function.

**2.4 Vectorized implementation of Moran's *I* in single-cell data**

The Moran's *I* measurement is also included in the esda package, but to fit the concept and speed up the computation in the single-cell scenario, we implemented it in our package with a vectorization strategy. The Moran's I of feature $\mathbf{x} \in \mathbb{R}^n$ in the single-cell data with *n* cells is defined as:

$$I_{\mathrm{x}} = \frac{\sum_i^n \sum_j^n w_{ij}(x_i - \bar{x})(x_j - \bar{x})}{\sum_i^n (x_i - \bar{x})} \cdot \frac{n}{\sum_i^n \sum_j^n w_{ij}}$$

where $w_{ij}$ is the connectivity weight between cell *i* and *j* in the KNN graph, and $\bar{x}$ is the mean of **x**. Suppose $\boldsymbol{W}$ is the row-standardized (row sums are all 1) connectivity matrix, **z** is an z-scored form of $\mathbf{x} \in \mathbb{R}^{n\times 1}$, Moran's *I* of **x** can be calculated with:

$$I_{\mathrm{x}} = \frac{\sum_i^n \sum_j^n w_{ij} z_i z_j}{\sum_i^n z_i^2} \cdot \frac{n}{\sum_i^n \sum_j^n w_{ij}}$$

$$= \frac{\sum_i^n \sum_j^n w_{ij} z_i z_j}{\sum_i^n z_i^2} = \frac{\mathrm{z}^T(\mathrm{Wz})}{\mathrm{z}^T \mathrm{z}}$$

In the single-cell scenarios, we need to calculate thousands of genes and peaks Moran's *I* values with the same connectivity matrix. To speed up, we vectorized the formula again to include all features:

$$\mathrm{I} = [I_1, I_2, \ldots, I_p] = \mathit{ColumnSum}(\mathrm{WZ} \circ \mathrm{Z}) \oslash \mathrm{ColumnSum}(\mathrm{Z} \circ \mathrm{Z})$$

where $\mathrm{Z} \in \mathbb{R}^{n \times p}$ is the feature matrix with *p* features, $\circ$ is the Hadamard multiplication, and $\oslash$ is the Hadamard division. We tested our implementation function, compared it with the result from the *esda* package, and ensured that they were the same.

**3. Regulatory marker discovery**

With the cluster annotation labels, users can find the differentially correlated gene-peak pairs in each cluster or compare clusters by simply calling the function *FindAllMarkers* or *FindMarkers*. The function will calculate cluster-specific local *L* matrix, make a statistic t-test comparison, summarize all the results, and return a data frame to the user containing mean correlations within the cluster, significance test results, and feature sparsity. To select the markers, users can call the function *MarkerFilter* to filter all pairs by feature sparsity, mean correlation difference, and p-values. If the plot parameter is set to True, a volcano plot will be returned together with the filtered data frame.

**4. Unlabeled regulatory state analysis**

When the cluster label is unavailable (usually when analyzing a single cluster), the regulatory state matrix can be utilized for various analyses, including sub-clustering, trajectory inferring, and feature module discovery.

The function *RegulatoryAnalysis* can cluster all the pseudo-bulks or cells, from which the trajectory will be inferred with *Scanpy* functions. This function can be repeatedly called for users to tune parameters like clustering resolution, and the root cluster for trajectory.

After determining the trajectory pseudo-timeline, users can call the function *Time_Module* to discover (gene or peaks) modules by self-organizing map (SOM) from the user-provided feature list. The SOM algorithm is an unsupervised artificial neural network that groups all observations in a low-dimensional representation while preserving the topological structure of the data. Here, we take genes or peaks as observations and the averaged expression levels within each pseudo-time bins as features, and clustered the genes or peaks by SOM. Users can run the function multiple times to have the optimized results by changing the SOM shape (how many modules are needed and what is the similarity among all modules), learning rate, number of iterations, and regulation power sigma. While optimizing the SOM results, users can visualize the results by calling the *visualize_module* function to intuitively understand the results.

**5. Functional analysis**

From the global *L* index, regulatory marker discovery, or regulatory state analysis results, users will end up with a list of gene-peak pairs of interest. To facilitate the understanding of these pairs, we implemented functions to call *Homer* from inside the Jupyter notebook or Python command line (users need to have *Homer* installed).

The function *run_HOMER_motif* will prepare the input from the list of peaks users provide, set up the directory where the results will be generated, call Homer, and save all the results.

With the motif of interest, the function *motif_summary* will interpret Homer's results, extract all peaks with the motif enriched, and return the summarized data frame to users. With the peaks extracted, genes linked to them can be easily found and sent for functional annotations. This is unavailable in Homer and quite trivial to do without the function.

#### Supplementary Figures

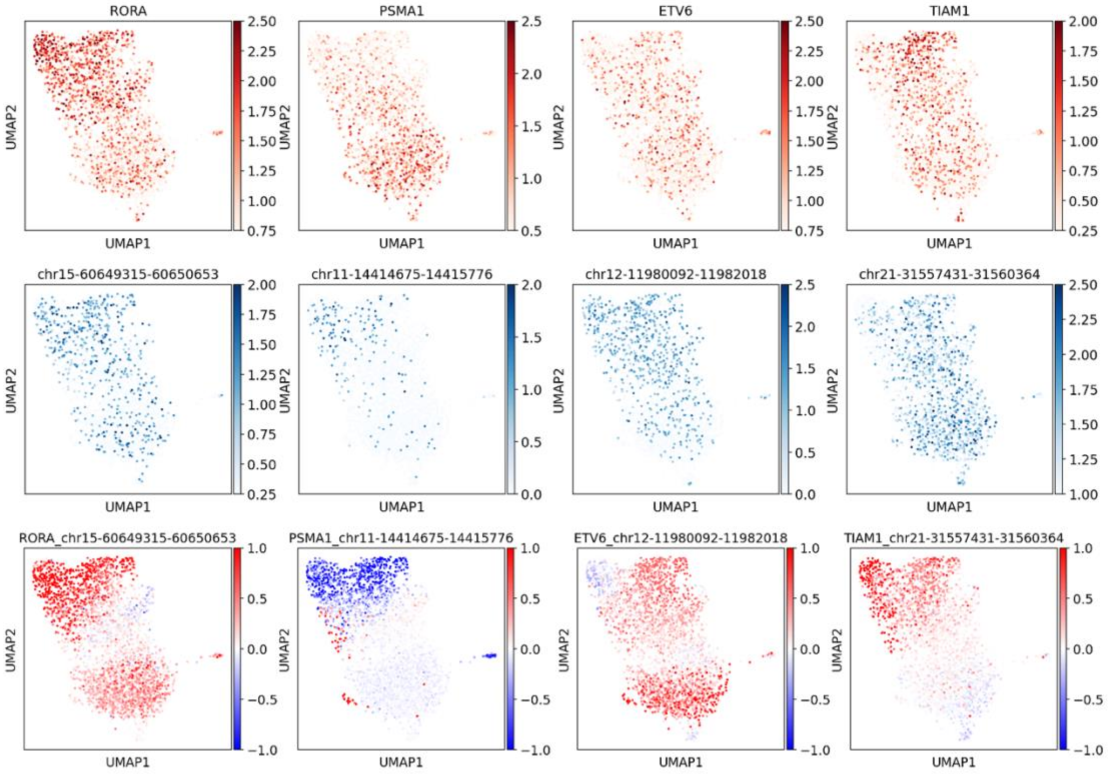


**Supplementary Figure 1.** Four examples show the heterogeneity of regulatory relationships within the CD4+ T cell cluster. Different gene-peak pairs can have different regulatory patterns.

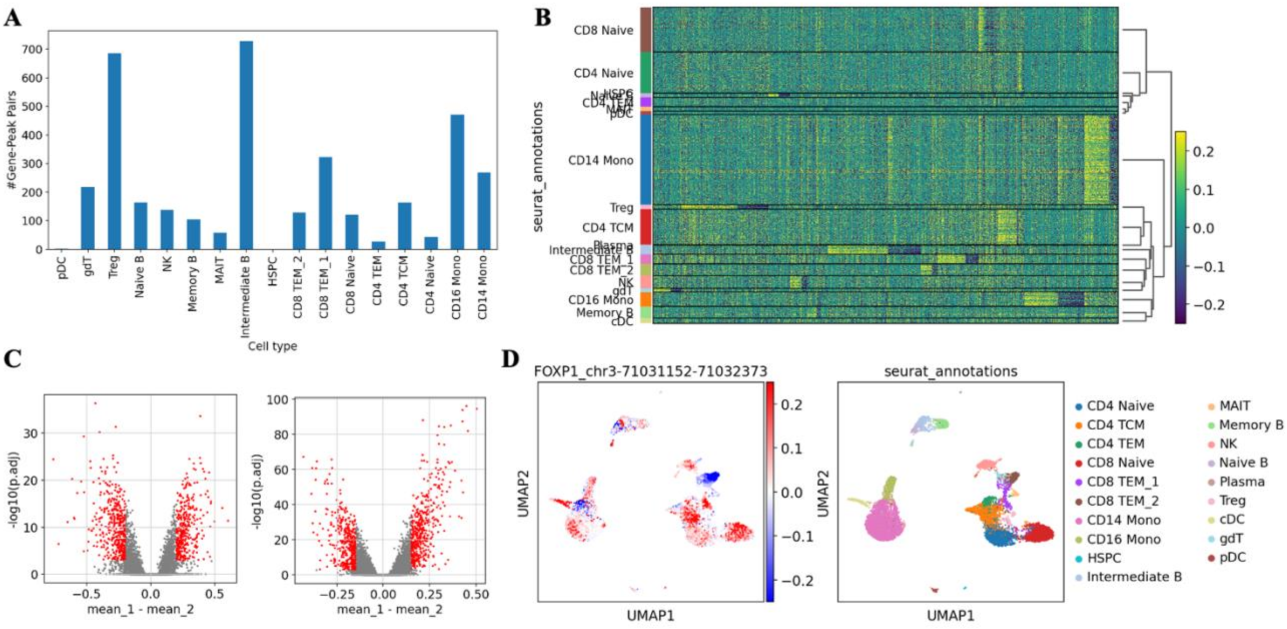


**Supplementary Figure 2.** (A) Bar chart summarizes the number of gene-peak markers found in each cell type. (B) Heatmap visualization of all gene-peak markers in pseudo-bulk data. Each row is a pseudo-bulk, each column is a gene-peak pair, and color indicates the correlation level. (C) Volcano plots showing the marker selected after filtering by mean correlation difference, adjusted p-value, and feature sparsity in intermediate B vs. memory B cells (left), and comparison among three CD4+ T cell subclusters (right). (D) Example showing the local *L* index can be mapped back from pseudo-bulk to single-cell data in scGREAT.

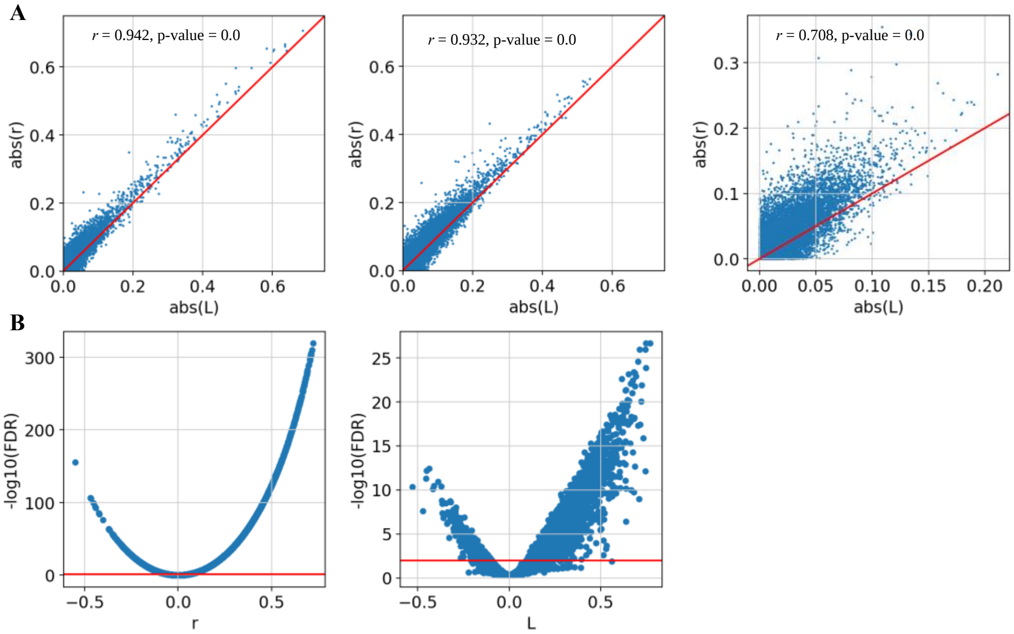


**Supplementary Figure 3.** (A) The consistency between L and r in studying the homogeneity of regulatory relationships was also seen in the cluster-specific analysis. Three clusters, including 10X Multiome PBMC CD4+ T cell, SHARE-seq mouse skin cluster 7, and SNARE-seq mouse brain cluster 2, were applied. (B) Distributions of p-values along the correlation level in Pearson's *r* results (left), and global *L* index results (right).

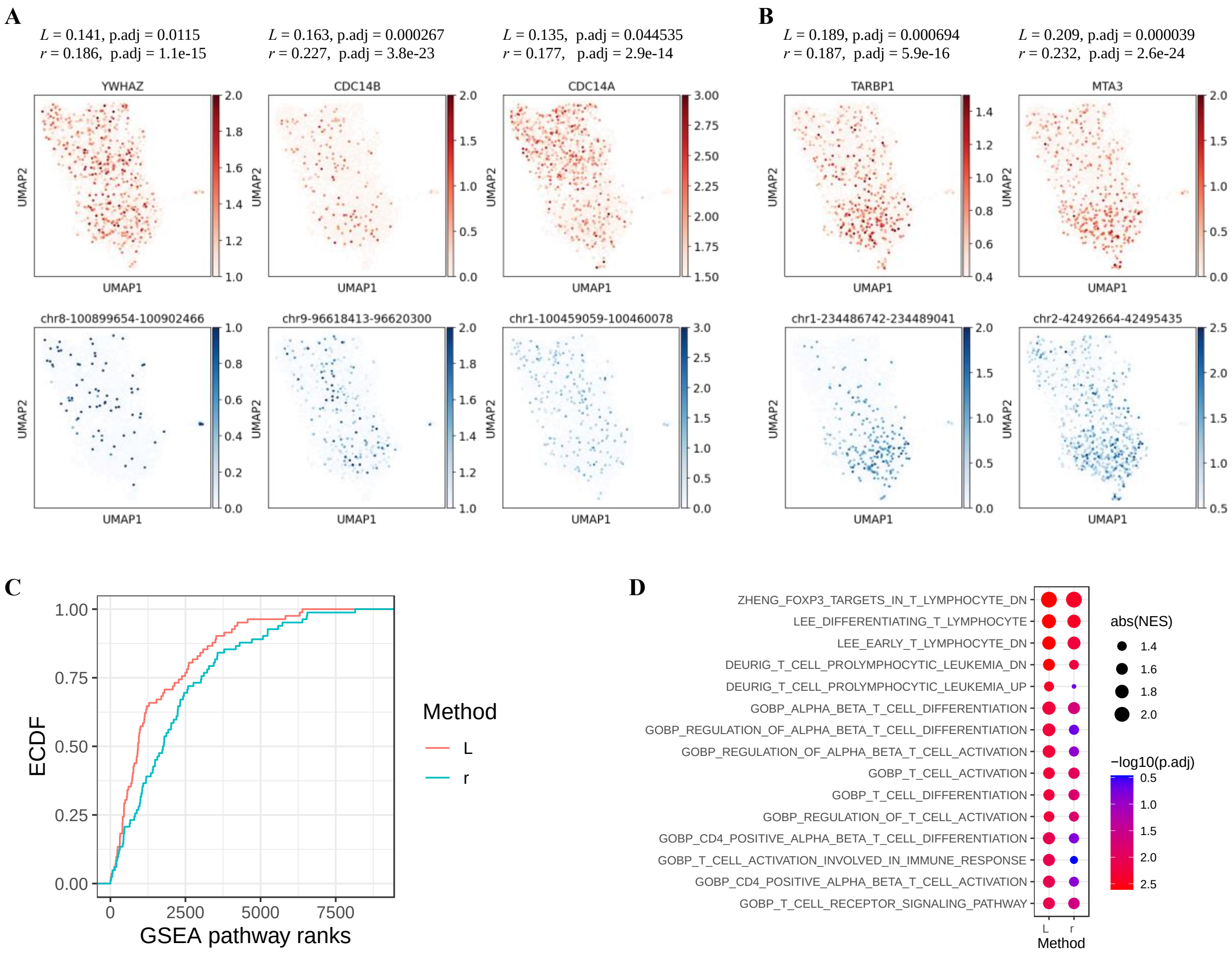


**Supplementary Figure 4.** (A) Feature plots show three examples of cell cycle genes and their nearby peaks. (B) Feature plots show two examples of immune-related genes and their nearby peaks with a similar Pearson's correlation to the cell cycle genes. (C) Empirical Cumulative Distribution of 89 T cell-related pathways' ranks. *L*-based ranks tend to give higher ranks for T cell-related pathways. (D) Gene Set Enrichment Analysis results using L-based and r-based gene ranks. Dot sizes represent the normalized effect size; color represents the -log10 adjusted p-value.
